# Machine learning prediction of resistance to sub-inhibitory antimicrobial concentrations from *Escherichia coli* genomes

**DOI:** 10.1101/2021.03.26.437296

**Authors:** Sam Benkwitz-Bedford, Martin Palm, Talip Yasir Demirtas, Ville Mustonen, Anne Farewell, Jonas Warringer, Danesh Moradigaravand, Leopold Parts

## Abstract

*Escherichia coli* is an important cause of bacterial infections worldwide, with multidrug resistant strains incurring substantial costs on human lives. Besides therapeutic concentrations of antimicrobials in healthcare settings, the presence of sub-inhibitory antimicrobial residues in the environment and in the clinics selects for antimicrobial resistance (AMR), but the underlying genetic repertoire is less well understood. We used machine-learning to predict the population doubling time and growth yield of 1,432 genetically diverse *E. coli* expanding under exposure to three sub-inhibitory concentrations of six classes of antimicrobials from single nucleotide genetic variants, accessory gene variation and the presence of known AMR genes. We could predict cell yields in the held-out test data with an average correlation (Spearman’s ρ) of 0.63 (0.32 - 0.90 across concentrations) and cell doubling time with an average correlation of 0.47 (0.32 - 0.74 across concentrations), with moderate increases in sample size unlikely to improve predictions further. This points to the remaining missing heritability of growth under antimicrobials exposure being explained by effects that are too rare or weak to be captured unless sample size is dramatically increased, or by effects other than those conferred by the presence of individual SNPs and genes. Predictions based on whole genome information were generally superior to those based only on known AMR genes, and also accurate for AMR resistance at therapeutic concentrations. We also pinpointed genes and SNPs determining the predicted growth and thereby recapitulated the known AMR determinants. Finally, we estimated the effect sizes of resistance genes across the entire collection of strains, disclosing growth effects for known resistance genes for each strain. Our results underscore the potential of predictive modelling of growth patterns from genomic data under sub-inhibitory concentrations of antimicrobials, although the remaining missing heritability poses an issue for achieving the accuracy and precision required for clinical use.

**Importance:** Predicting bacterial growth from genome sequences is important not only for a rapid characterization of strains in clinical diagnostic applications but for the identification of novel targets for drug discovery. Previous studies examined the relationship between bacterial growth and genotype in mutant libraries for laboratory strains, yet no study has so far examined the prediction power of genome sequences for bacterial growth in natural strains. In this study, we used a high throughput phenotypic assay to measure bacterial growth of a systematic collection of natural *Escherichia coli* strains and then employed machine learning models to predict bacterial growth from genomic data under non-therapeutic sub-inhibitory concentrations of antimicrobials that are common in nonclinical settings. Our results revealed a moderate to strong correlation between predicted and actual values for different antimicrobials concentrations. Furthermore, the quantified effect of resistance genes on bacterial growth indicate these genes are still effective at sublethal antimicrobial concentrations.

## Introduction

*Escherichia coli* is a dominant bacterial species in the lower intestine of humans and other endotherms, as well as within a range of environmental niches (1). Over recent decades, the rising frequencies of *E. coli* that are resistant to multiple antimicrobials in both the clinic and the environment have become a source of serious concern for human and live-stock health (2). A remarkably broad and flexible genetic repertoire spanning both the core and the accessory genome appears to underlie this rapid spread of multiresistant *E. coli*. To diagnose and understand this multiresistance, we must accurately and exhaustively estimate how the variants in this genetic repertoire, individually and in combination, affect *E. coli* growth under exposure to the range of antimicrobial concentrations that the species encounters in nature and in the clinic.

The use of the *E. coli* gene knockout (KO) library captures the effect of complete loss of many individual genes in the K-12 genome on growth in sub-lethal antimicrobial concentrations (3, 4). Combined with mechanistic modelling, such data can guide our understanding of some of the K-12 AMR defence systems. However, natural *E. coli* have an open pan-genome, which results in a high level of population diversity (5). Thus, the genome of K12 may not represent the diversity, given the phylogroup of K-12 only harbours ~20% of genes across the pan-genome for known *E. coli* lineages and a similar fraction of the single nucleotide diversity present in the species (6, 7). As a result, most AMR effects likely evade detection in K-12 screens while those that are detected often have little or no effect in other backgrounds. Genome-wide association studies (GWAS) on broader panels of clinical and environmental *E. coli* lineages could successfully estimate moderate or large AMR effects of common variants of all types (8). However, rare, weak and background dependent AMR effects are challenging to detect and measure. Linkage also means that some detected variants, although they may serve as diagnostic biomarkers for AMR, provide little molecular understanding of AMR itself. Moreover, these methods examine the association of the presence/absence of variants with the phenotype in isolation from the rest of variants, thus neglecting potential phenotypic interactions between variants (9).

Machine-learning methods incorporate genomic variants into a single prediction framework that can capture the background dependency of AMR effects. Over recent years, the number of machine learning models for predicting AMR from whole genome sequencing data has risen sharply (10–12). Ensemble models that combine results from multiple weak learners into one model proved superior to other models (13). So far, most studies have focused on qualitative AMR data sets with lineages classified as resistant or sensitive to diagnostic or minimum inhibitory (MIC) antimicrobial concentrations and models are judged based on how accurately this binary classification is recalled, meaning that much of the underlying biological variation has been obscured (reviewed in (11)). As a result, published approaches have mostly captured common and large AMR effects, which represent already well understood aspects of AMR biology. However, the impact of rare, weaker variants as background dependent contributions on bacterial growth under AMR treatment has not been examined (11). Furthermore, the evolution of antimicrobial resistance in nature is likely driven by antimicrobial concentrations that are much lower (< one hundredth) than the MIC or the diagnostic concentrations that are used in the clinic (14). Antimicrobials can reach high local concentrations downstream production plants and sewer outlets (15), but are diluted to a 100-fold below MIC in many environmental and wild animal niches where *E. coli* is more frequent (14, 16, 17). It is conceivable that the natural selection for AMR resistance at sub-MIC concentrations may be driven largely by effects other than those controlling resistance at diagnostic concentrations.

We here used machine-learning to predict the population doubling time and cell yield of >1,400 *E. coli* quantitatively at sub-MIC antimicrobial concentrations from genomic data with no prior information on resistance mechanism. We quantified the contribution of both known AMR determinants and genetic variants previously not known to affect AMR, disclosing the importance of the processes cell wall biosynthesis processes and for carbohydrate metabolism. Despite the unprecedented scale of the study and the low measurement error, the best model predictions were limited to an average correlation (Spearman’s ρ) of 0.63 (ranging from 0.32 to 0.90 across antimicrobials) for cell yield and 0.47 (ranging from 0.32 to 0.74 across antimicrobials) for doubling time across antimicrobials. The unaccounted heritability of antimicrobials resistance is therefore substantial, underscoring the challenge of fully explaining AMR in an enormously diverse bacteria where most causal variants are rare or only rarely affect AMR.

## Methods

### Strains and genomic data

We used data for 1,432 *E. coli* lineages which were collected, sequenced and extensively growth phenotyped in the ongoing TransPred project and are available online as part of an expanding resource (http://www.github.com/matdechiara/TransPred). The here included strains were recovered from diverse human settings, including hospital and community onset infections, food, wild animals and wastewater treatment plants. All TransPred sequencing was performed at the Wellcome Sanger Institute with a 450-bp insert size on Illumina HiSeq2500 machines with paired-end reads with a length of 100bp. The sequence data for the here included stains is submitted to the European Nucleotide Archive (ENA) under the study accession number of PRJEB23294. For the purpose of predicting phenotypes from genomic information, we here assembled the paired-end reads of these strains using an assembly and improvement pipeline (18) based on Velvet (19). These *de novo* assemblies were annotated with Prokka (20). Codes and other intermediate files are available in http://www.github.com/dmoradigaravand/TransPred_ML.

### Genomic and phylogenetic analysis

For the purpose of identifying genomic variants that are used by models for predictions of growth at sub-inhibitory antimicrobials concentrations, we here mapped the short reads against the K-12 *E. coli* reference genome (accession number: NZ_CP032667) with SMALT v 0.7.4 (https://www.sanger.ac.uk/resources/software/smalt/) (21). A threshold of 30 was used for mapping and SNPs were subsequently called and annotated SAMtools mpileup (22) and BCFtools (23). We removed SNPs at heterogeneous mapping sites in which the SNP was present in less than 75% of the reads at the site.

To reconstruct the alignment free phylogenetic tree, we first enumerated *k*-mers of size 50 in the genome and then counted the number of *k*-mers shared between pairs of isolates. The resulting a similarity matrix was subsequently converted into a distance matrix, which was used to create a neighbour-joining tree with the Ape package in R (24).

To reconstruct the pan-genome of the whole collection, we fed the output of Prokka (20) into Roary (25) and used the identity threshold of 95% to identify gene orthologous families. For functional enrichment analysis, COG categories of genes were extracted from the annotation by Prokka and assigned to functional classes, as defined in (26). To fully annotate resistance gene families, e.g. tetracycline resistance efflux genes, we conducted blast for the genes used for prediction by the models on the CARD database. We further explored the functions of hit genes by performing blastx using NCBI protein database.

We utilized Scoary (27) with 50 permutations to compute the association between predictive accessory genes that were identified by the model and continuous growth features, i.e. cell doubling time and growth yield, at each antimicrobial concentrations. Since Scoary works on binary response features, we first binarized the continuous response variables according to the median value for each growth feature. We used the worst pairwise computed *p-*value in Scoary, which reports the significance of association of the variant with population structure, as the *p*-value. Antimicrobial resistance determinants were identified using ARIBA with the default similarity threshold (16). The output of ARIBA was then turned into a presence/absence matrix.

### Growth at sub-MIC antimicrobial concentrations

We used the population doubling time and cell yield data generated for the included strains in the TransPred project (http://www.github.com/matdechiara/TransPred). The data captures the population doubling time and growth yield after 8 hours of growth of 1432 *E. coli* lineages expanding clonally as individual colonies on top of a solid matrix composed of LB medium in which either of three sublethal concentrations for six antimicrobials, CIP (ciprofloxacin), CTX (cefotaxime), TET (tetracycline), TRIM (trimethoprim) and CAM (chloramphenicol), had been embedded. Colonies were deposited as initially isogenic populations at initial population sizes of ~100,000 cells, with 1536 colonies deposited in systematic colony arrays on each plate using robotics. 384 of these colonies were identical controls used to correct for spatial bias between and within plates. For each concentration of each antimicrobial, each lineage was cultivated as six biological replicates on different plates. Population expansion for each colony was followed by measuring cell numbers at 10 minute intervals using the Scan-o-matic framework, version 2.0 (28). From each colony growth curve, the cell doubling-time, and the total cell yield after 8h was extracted (growth yield). Experiments included automated transmissive scanning and signal calibration in 10 min intervals, as described (28). The absolute population doubling times and yields were log(2) transformed and normalized to the corresponding measures of adjacent controls on each plate, while data for missing or mis-quantified colonies were discarded, and the median of these logged and normalized values across biological replicates for each lineage was retained and used as the response continuous variable in the machine-learning setting.

### Machine learning to predict growth at subMIC antimicrobial concentrations from genome

We calculated features from the genome to use as predictors in the models. These predictors included: 1) pan-genome gene presence: for each accessory gene, and for each *E.coli* strain, a binary indicator of gene presence in the strain as determine by Roary 2) SNPs: which included distinct encodings for alternative nucleotides and for the absence of the site as compared with the reference genome and 3) resistome: for each strain, for each of the resistance determinants from the CARD database (17), a binary indicator of their presence in the strain.

We applied three classes of machine learning methods on predictor matrices. These consisted of a lasso regularized regression model (29), a gradient boosting regressor ensemble model (30) and a feed forward residual neural network (31). The models were trained on 80% of the input dataset, with two-fold cross-validation for tuning hyper-parameters, and evaluated on a 20% random held-out dataset. We excluded rows with missing values for each drug. For the lasso regression and gradient boosted regressor models, we utilized the Sklearn package (32) and the functions of <u>linear_model</u>.Lasso and Ensemble.gradientBoostingRegressor, respectively. We used mean absolute estimates for error, which compared to other error measures of statistical dispersion better accounts the impact of outliers, to choose the best performing model. The strength of the correlation between the predicted and real data was measured using the Spearman’s rank correlation coefficient (Spearman’s ρ).

We used a grid search to find optimum values for the hyperparameters. The lasso model was tuned by finding the value for the penalty term, testing values [10^−5^, 10^−4’^, 10^−3^, 10^−2^, 10^−1^]. We tuned the gradient boosted regressors by finding the optimal values for key tree related parameters, including tree depth [10, 30, 50] and number of iterations [100, 150, 200] using a grid search method. Manual inspection of error in runs over iterations revealed that models began to overfit after generations 100 for the response variable.

To develop the neural network, we used the Keras library (33), and trained fully connected feed-forward networks with drop-out of 0.2. We split the input data test (20%), validation (20%) and training (60%) datasets, and tested *n*=2, 4 or 6 hidden layers, and n=20, 40 or 60 number of neurons per layer, which were screen jointly. The Adam algorithm (34) was used for an adaptive learning rate optimization. Models were trained for at most 100 epochs, with batch size of 32.

We employed the EarlyStopping feature in the Keras library to terminate the training as soon as the validation loss worsened from one epoch to the next. Moreover, we examined the inclusion of a skip connection feature in the model, which skips some layer in the neural network and feeds the output of the first layer as the input to the last layer. The skip connection has been suggested to mitigate the problem of vanishing gradients, and therefore, speed up the learning process (31). We reported the model with the highest values for the Spearman’s ρ between the predicted and actual value in the held-out data as the best performing neural network model.

To account for the effect of measurement errors on prediction accuracy, we randomly drew one measurement from six replicas for each strain in the test dataset to create an alternative bootstrapped test dataset. We then repeated the process to create 100 alternative test datasets for each condition. We then computed Spearman’s ρ between predicted values and bootstrapped test data in each of the 100 bootstrapped test datasets. We subtracted the mean of 100 Spearman’s ρ values from one to quantify the effect of variation in measurements on prediction for each condition. We amended prediction performance values after correcting for the error by dividing the correlation coefficient from predictions by the mean correlation coefficient, computed for randomly generated alternative test datasets.

### Prediction of resistance and susceptibility labels at therapeutic concentrations

We examined whether the population doubling time and growth yield predicted at sub-MIC antimicrobial concentrations well reflected antimicrobial resistance at near diagnostic MIC concentrations. To do this, we used the resistant/susceptible labels against CTX, CIP and KAN for 273 of our strains, which we reported in a previous study (35). In that study, we had empirically measured whether each strain was capable of growing at near diagnostic MIC concentrations and labelled them accordingly as resistant and susceptible at therapeutic concentrations of antimicrobials. We then trained the gradient boosted regressor models on a training/validation dataset, excluding the 273 strains, and used the tuned model to predict cell yield and doubling time for the 273 strains with known phenotypic labels, i.e. resistant and susceptible. Distribution of growth-related features was bimodal, corresponding to resistant and susceptible strains, with modes separated more as the concentrations of antimicrobials increase (Figure S4A). To convert continuous predicted values into binary values, we first fitted a bimodal distribution on predicted values with Expectation Maximum algorithm implemented in the normalmixEM function in the mixtools R library (36). We then used the posterior distributions for the fitted distributions to assign resistance and susceptible labels, assuming the distribution with smaller mean corresponds to the susceptible sub-population, as shown in an example in Figure S4B. Therefore, labels were assigned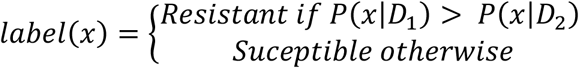, where *D*_1_ and *D*_2_ are parameters of the normal distributions of the two mixture components and 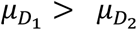. We then measured precision (fraction of true positive instances among the retrieved instances), and recall (fraction of retrieved true positive instances among the true positive instances), to assess the performance of classification.

### Extracting genomic features important to growth at subMIC antimicrobial concentrations

We adopted global and local approaches to compute the importance of genomic features for predictions in the gradient boosted regressor tree models. In the global approach, we used the feature importance calculator (feature_importances) as part of the Sklearn learn package, where genomic feature importance was computed during the optimisation of the weak learner in the boosting process. Here, the importance value for a genomic feature corresponds to the fraction of samples for which the tree will traverse a node that splits based on the feature. These values were averaged across all the trees during the iterations. In order to identify the features that most robustly contributed to prediction, we repeated the prediction on the test data set with the tuned model for 20 times, computed the frequency at which each genomic feature occurred and then averaged rank and importance for each feature across the 20 replicates.

The local explanation approach was post-hoc and employed the game theoretic concept of SHAP (SHapley Additive exPlanations), derived from coalitional game theory, to explain the output of machine learning models for each sample (37). The SHAP explanation method is an additive feature attribution method. Here, predictor feature values of data points serve as players in a coalition. A player can either be an individual genomic feature value or a group of feature values. The Shapley value of a feature value is defined as the contribution of the predictor to the payout, which is weighted and integrated over all possible feature value combinations. Shapley values are computed by introducing each feature, one at a time, into a conditional expectation function of the model’s output, *f*_*x*_(S)= E⟨*f*(X)|*do*(X_*s*_ = *x*_*s*_)⟩ and attributing the change produced at each step to the feature that was introduced; then averaging this process over all possible feature orderings. Here S denotes the set of features we are conditioning on, X is a random variable representing the model’s input features, *x* is the model’s input vector for the current prediction. We used the TreeExplainer function to explain the fitted tree as implemented in (38) available in http://www.github.com/slundberg/shap).

Besides single predictors, the method allows to determine the mean combined effect for multiple predictors. We obtained such combined effects of specific antimicrobial genes for TRIM on growth yield by measuring the mean values of SHAP values for single resistance genes in different contexts resulting from combinatorial patterns for the presence and absence of other resistance genes. The combinations included four possible cases for the presence and absence of TRIM genes, i.e. *dfrI* and *dfrV*, which was used to reconstruct a fitness landscape.

To assess the prediction importance of SNPs, we first extracted the global importance values for predictors consistently used by the gradient boosting regressor model, as detailed above. Since we included SNPs in both core and accessory genome, we specifically tested whether the presence of a base-pair substitution at the site -and not the presence/absence of the site- accounts for the importance of the feature in the predictive model. To this end, we conducted an ANOVA test to examine the significance of the association between the presence of each nucleotide substitution and the growth-related dependent variable. We repeated the process for every base alteration for SNP sites that were found important by the gradient boosting model. Furthermore, we identified non-synonymous SNP sites in the list of important SNPs that involved more than one type of nucleotide alteration (multivariate sites). The independent emergence of different genetic variants across lineages at these sites is suggestive of the adaptive significance the SNP and/or the gene, where the SNP occurs.

### Pan-genome simulation

To assess the impact of pan-genome and population parameters on prediction, we simulated pan-genome evolution under different combinations of the evolutionary constraints of gene-gain/gene-loss ratio and population size. We then used the pan-genome data to predict the simulated cell yields, with increasing values for the penetrance of a causal allele. We simulated pan-genomes using the Simurg package in R (39). To simulate different evolutionary scenarios and trait distributions, we first used a range of values for population size (N:200, 400 and 600) and the rate of gene acquisition (*υ*: 10^−7^, 10^−8^ and 10^−9^), which tunes the pan-genome size, to create pan-genome matrices. We kept the rate of gene loss (γ) at a constant value of 10^−11^. This resulted in pan-genomes with varying sizes. We then randomly drew an accessory gene with a frequency in the range of 0.45-0.55 and labelled them as causal. We assigned continuous growth values to strains using a distribution after fitting a normal curve with the mean values of μ and standard deviation of σ to a realistic distribution, i.e. the distribution of growth yield values in the absence of drug treatment (Figure S4A). For strains lacking the causal gene, growth values were randomly drawn from the baseline normal distribution with a mean value of μ, whereas for strains harboring the causal gene, random growth values were drawn from a normal distribution that had a mean value of μ+λσ, where λ corresponds to the selective advantage of the resistance gene. We screened a wide range of values for λ (0, 0.05, 0.1, 0.25, 0.5, 1, 1.5, 2, 5 and 10). We tuned and tested gradient booting regressors on the predictor and dependent datasets, as described above.

In order to compare simulation and actual results, we estimated the value for λ under antimicrobial treatments. To this end, we first fitted a bimodal mixed distribution to growth yield and doubling time distributions for each condition (Figure S4A), as mentioned above, and then measured the difference between the means of normal distributions for susceptible and resistance sub-populations, as the experimental λ for the condition.

## Results

To predict *E. coli* growth at subMIC concentrations of antimicrobials, we used >1400 clinical, commensal, and environmental *E. coli* strains from major globally circulating lineages in the TransPred project (http://www.github.com/matdechiara/TransPred). We used available whole genome sequence information and data on the population doubling time and growth yield of each isolate when clonally expanding as observations in our prediction framework. We selected three subMIC concentrations of six bacteriostatic and bactericidal antimicrobials effective against *E. coli* as measurement contexts.

In *E. coli*, resistance to diagnostic concentrations of the antimicrobials used here occurs mainly through the horizontal transfer of plasmid-borne accessory genes that vary in presence/absence across lineages. We therefore first probed how well we could predict growth at subMIC antimicrobial concentrations from pan-genome data, using 35,641 genomic features with unique presence-absence distributions of gene families across our lineages, and linear, ensemble and neural network regressors. Based on gene presence-absence, our predictive models attained a mean of 0.52 (range: 0.25, 0.73) and 0.40 (range: 0.28, 0.61) of Spearman’s *ρ* for growth yield and doubling time, respectively, in held-out test data (Figure 1). Because the prediction error rates for the training and test datasets were comparable (Figure S1), models were not overfitted and unlikely to have been negatively affected by the number of predictor features vastly exceeding the sample size. The correlations between predicted and actual values were all significant, when compared to the median correlation value computed for randomized datasets (Figure S2). The observed improvement in Spearman’s *ρ* values when we compared predicted value with randomized values were on average 0.37 (range: 0.27-0.64) for population doubling time and 0.47 (range: 0.20-0.74) for growth yield (Figure S2). In 8 out of 12 out of 18 conditions with antimicrobial treatment the measures improved with increasing antimicrobial concentrations, likely due to the higher fraction of between strain variation explained by genetics, i.e. a higher broad sense heritability, at these concentrations (Figure 1). Amongst predictive models, the gradient boosting regressor outperformed lasso and neural networks for 30/38 conditions, underscoring the suitability of these models for genome-based prediction of AMR (10, 35, 40). In 7/38 conditions, the neural networks were superior.

**Figure 1.**
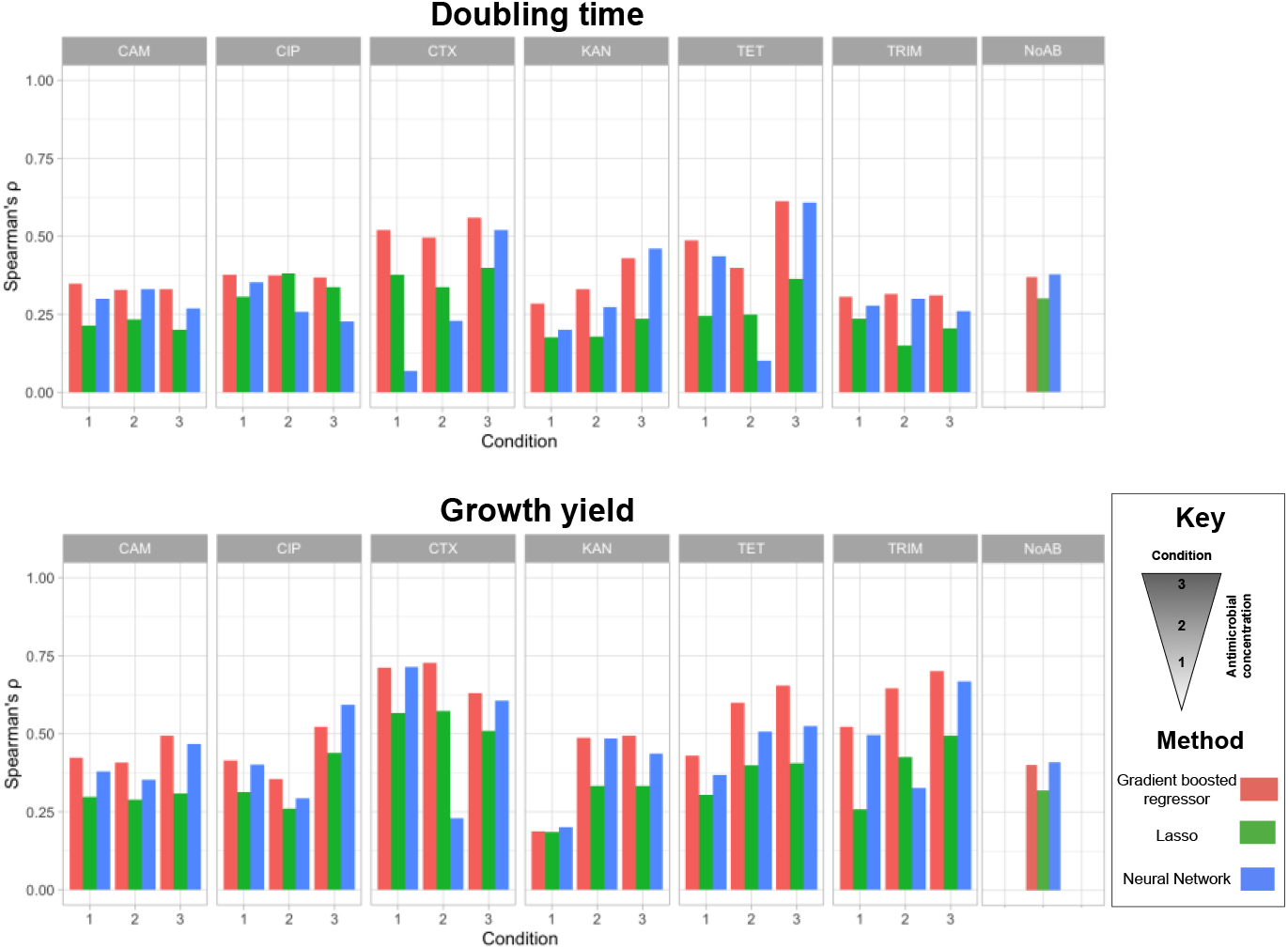
The performance (Spearman’s rho, y-axis) of three predictive models (colors) for 6 antimicrobials (panels, x-axis) under 3 concentrations (x-axis) and a control condition for doubling time (top row) and growth yield (bottom row). The performance was assessed as the magnitude of correlation between the predicted and real data in the test dataset. Numbers 1, 2,3 represent the increasing MIC-subinhibitory concentrations.

Despite a large data set, even the best performing model failed to explain on average 0.54 of the variance across conditions when only taking accessory gene variation into account. This was not due to environmental variation or stochastic noise, because measurements’ errors were overall low, with average Spearman’s *ρ* between bootstrapped replicates of 0.88 (range:0.85, 0.91, see Methods). Thus, most of the unexplained variation might be genetic in nature and reflects missing heritability. Conceivably, accessory genes that are too rare, too weak or too dependent on the presence of other variants that are rare to be captured by our models, could account for some of this missing heritability. Such genetic effects stand a larger chance of being captured if the sample size is increased. We therefore examined whether moderate variations in sample size substantially impact on prediction accuracy by down-sampling the included strains. We trained the models on randomly generated sub-samples of our dataset at different sizes and examined the performance of the trained on the same held-out test dataset. Figure S3 reveals that for the majority of conditions the performance of the models tends to level out with increasing training sample size. In the absence of antimicrobial treatment, sub-samples that were half of the full training set produced similar results to predictions with the full training set (Figure S3A and S3B). We only detected monotonic improvement with increasing sample size for 4/18 (population doubling time) and 1/18 (growth yield) conditions (Figure S3A and S3B). These findings show that predictions are unlikely to improve substantially with moderate increases in sample size, and that the missing heritability is either accounted for by accessory genes whose effects can only be captured by vast increases in sample size, or by genetic effects other than those associated to accessory genes (see Discussion).

We pursued the latter hypothesis by next asking to what extent predictions of growth at subMIC antimicrobial concentrations improved by including the presence and absence of individual SNPs in the core and non-core *E. coli* genome. We called 1,324,765 SNPs and included these in training dataset for training gradient boosting models (Figure 2). The inclusion of SNP information improved the pan-genome based prediction for 13/19 and 10/19 conditions for population doubling time and growth yield, respectively. The average prediction improvement for pan-genome based prediction was 6.1% (range:0.5%-17%) for population doubling time and 6.4% (range:0.6%-14%) for growth yield. Because SNP input matrix retains information on the presence and absence of genes, as we have examined SNPs in accessory genome as well, the prediction accuracy of models only based on SNPs could not be strictly evaluated. We conclude that SNPs whose effects are sufficiently common and strong to be captured by modelling on sample sizes of >1400 genotypes only accounts for a small fraction of the missing heritability of AMR.

**Figure 2.**
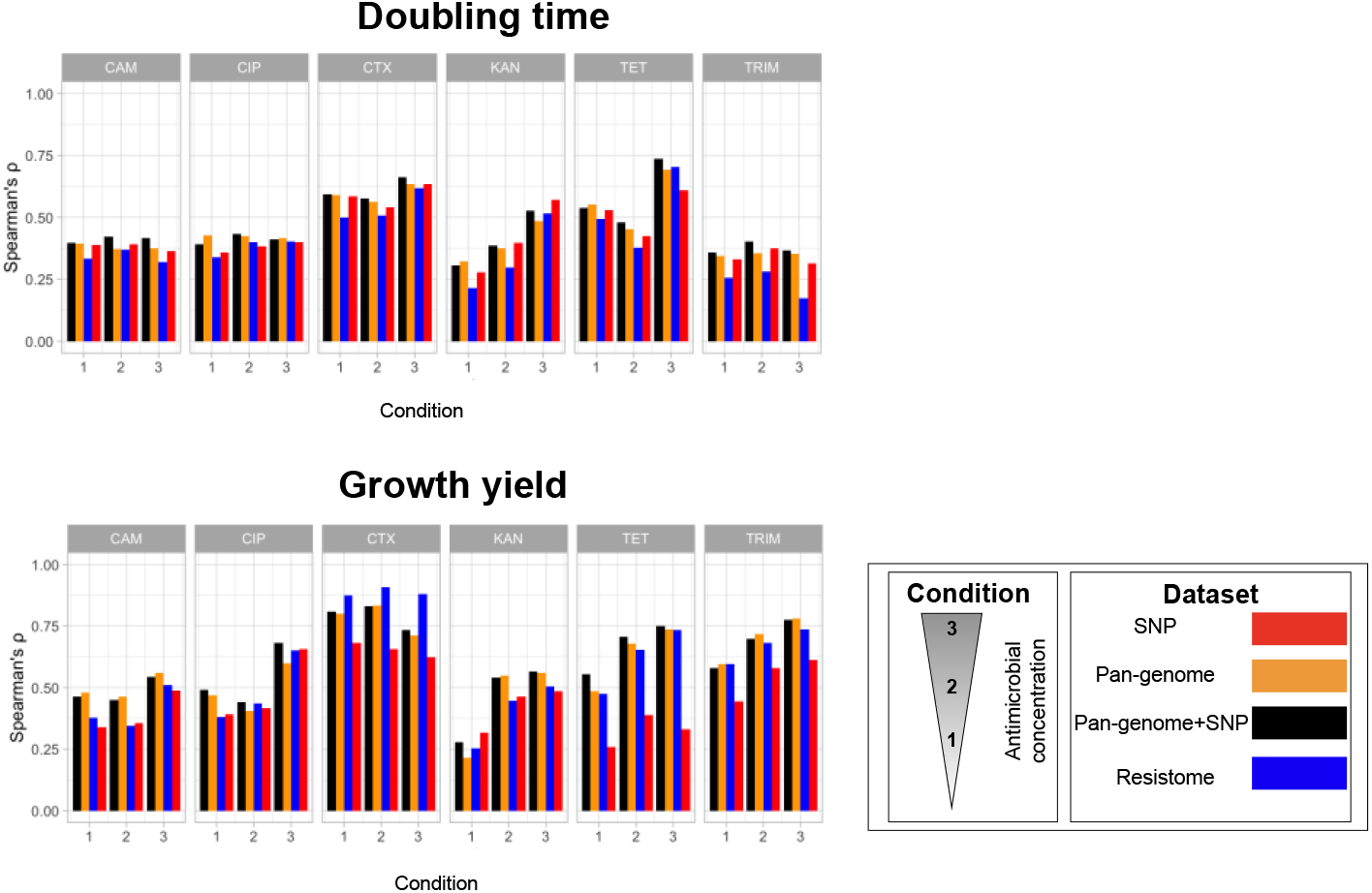
The prediction results (y-axis) measured as the correlation of real and predicted growth rate and doubling time values in the test dataset across antimicrobials and different subinhibitory concentrations (x-axis), using alternative feature sets (colors). Gradient regressor models were trained on resistome SNP, pangenome and combined SNP and pan-genome data. We have corrected for the effect of the variation in measurements.

The current standard for predicting antimicrobial resistance at diagnostic concentrations rely on detecting the presence of known resistance genes, together called the resistome. We therefore evaluated how well models exclusively based on the known resistome performed, as compared to models that use more complete genome information. Overall, we found predictions of models using only the resistome data to improve for both population doubling time and growth yield with increasing concentrations, with the prediction accuracy increasing monotonically with increasing antimicrobial concentrations for four (growth yield) and five (population doubling time) out of the six antimicrobials (Figure 2). The resistome also better predicted growth yield than population doubling time in 15/18 antimicrobial concentrations, suggesting that antimicrobial resistance genes generally enhance the cell yield more than they reduce the population doubling time (Figure 2). Growth yield values have a broader range, in comparison to doubling time, resulting in a stronger signal to noise values and a better performance of predictive models. The prediction performance worsens when we used the resistome only in that our models achieved a prediction performance that averaged 85% (range:46%-99%) of what the best performing models achieved using the complete genome information, i.e. considering all SNPs and gene presence-absence variations (Figure 2). Moreover, we achieved better model performance when exclusively using known resistance genes only for growth yields in CTX treatments, and this likely reflects the outsized effect of beta-lactamases, especially ESBL genes, on CTX resistance. We conclude that *E. coli* growth at sub-MIC antimicrobial concentrations is generally predicted best from whole genome data. Altogether, the best combination of predictive models and predictor datasets yielded Spearman’s *ρ* values with an average of 0.47 (range: 0.32- 0.74) for population doubling time and 0.63 (range: 0.32-0.90) for growth yields after accounting for the effect of measurement errors (Figure 2).

Antimicrobial resistance in clinical contexts is diagnosed as growth or non-growth at high MIC concentrations. To explore to what extent antimicrobial resistance at subMIC and MIC concentrations depends on the same genomic features, we first labelled our strains as resistant or sensitive, based on the distributions of predicted cell doubling times and growth yields (Figure S4A, S4B) (see Methods). We next measured how well these labels recalled and predicted those assigned for the same strains grown at higher MIC concentration. Overall, resistance-sensitivity labels assigned based on growth yield at sub-MIC concentrations assigned labels well captures those assigned at MIC concentrations, with average recall rates of 0.79 (range:0.58 (for CIP), 0.94 (for TRIM)) and an average precision of 0.77 (range:0.29 (for CIP), 0.94 (for CTX)) (Figure S4C). Labels assigned based on population doubling times at MIC concentrations less well captured labels at higher concentrations, with an average recall of 0.53 (range:0.07 (for CIP), 0.87 (for CTX)) and an average precision of 0.45 (range:0.10 (for CIP),0.83 (for CTX)) (Figure S4C). Resistance to diagnostic concentrations of CTX and TRIM, which is known to be mainly determined by the presence of particular accessory resistance genes, were better predicted by inferred growth measures at sub-MIC concentrations than resistance to CIP, which is held to be primarily driven by chromosomal mutations in core *E. coli* genes (Figure S4C). We compared the performance of best performing prediction models at sub-inhibitory MIC with previous machine learning models that were trained on large-scale pan-genomes and binary phenotypic labels (resistance vs susceptible) (35). Sub-inhibitory MIC based models attained 95% of reported precision and 99% of reported recall for TRIM and 95% of reported precision and 99% of reported recall for CTX, from the best performing predictive models in (35). This shows that in general *E. coli* growth predicted at subMIC antimicrobial concentrations well captures AMR phenotypes at diagnostic concentrations.

We next sought to understand which gene presence and absence variation contributed to predicting *E. coli* growth (see Methods) and first identified those important to population doubling time and growth yield in the absence of antimicrobials. We found 46 (growth yield) and 37 (population doubling time) genomic features (individual and sets of gene presence-absence variations) that the gradient boosting models consistently used (Figure 3A, 3B). SHAP plots confirmed that the presence of 27/46 and 22/37 of these features associated with better growth (Figure 3C, 3D). Only two of these genomic features were shared between growth yield and doubling time, consistent with these being genetically distinct aspects of *E. coli* growth. Most of the detected genes lacked functional classifications, but those with at least one functional annotation encoded membrane proteins (functional group M) were overrepresented (13% of genes compared to 2% in the whole pan-genome) (Figure 3A). Only a few features (2/37 for cell doubling time and 3/46 for cell yield) cooccurred across lineages, i.e. Pearson’s r > 0.90 for their associations with each other), excluding an extensive effect of the linkage between genetic features (Figure 3A, 3B). The majority of identified genes, i.e. 44/46 and 35/37 of features for growth yield and doubling time, respectively, were also distributed across clades in a manner that are associated the population structure. Among features with such phylogenetically distinct signals, i.e. not linked with population structure, two stood out. The simultaneous presence of the phage-like gene for an uncharacterized protein *yjhV* and *fecC*, which encodes an integral membrane protein component of the iron transporter in the citrate-dependent iron system (41, 42), associated strongly to population doubling time, in a manner that could not be explained by population structure. Similarly, the shared presence of *kpsD* and *rfbD* predicted a high growth yield. *kpsD* is responsible for the capsular polysaccharide translocation channel for cell wall synthesis, while *rfbD* encodes one of the precursors for the O-antigen of lipopolysaccharide (28).

**Figure 3.**
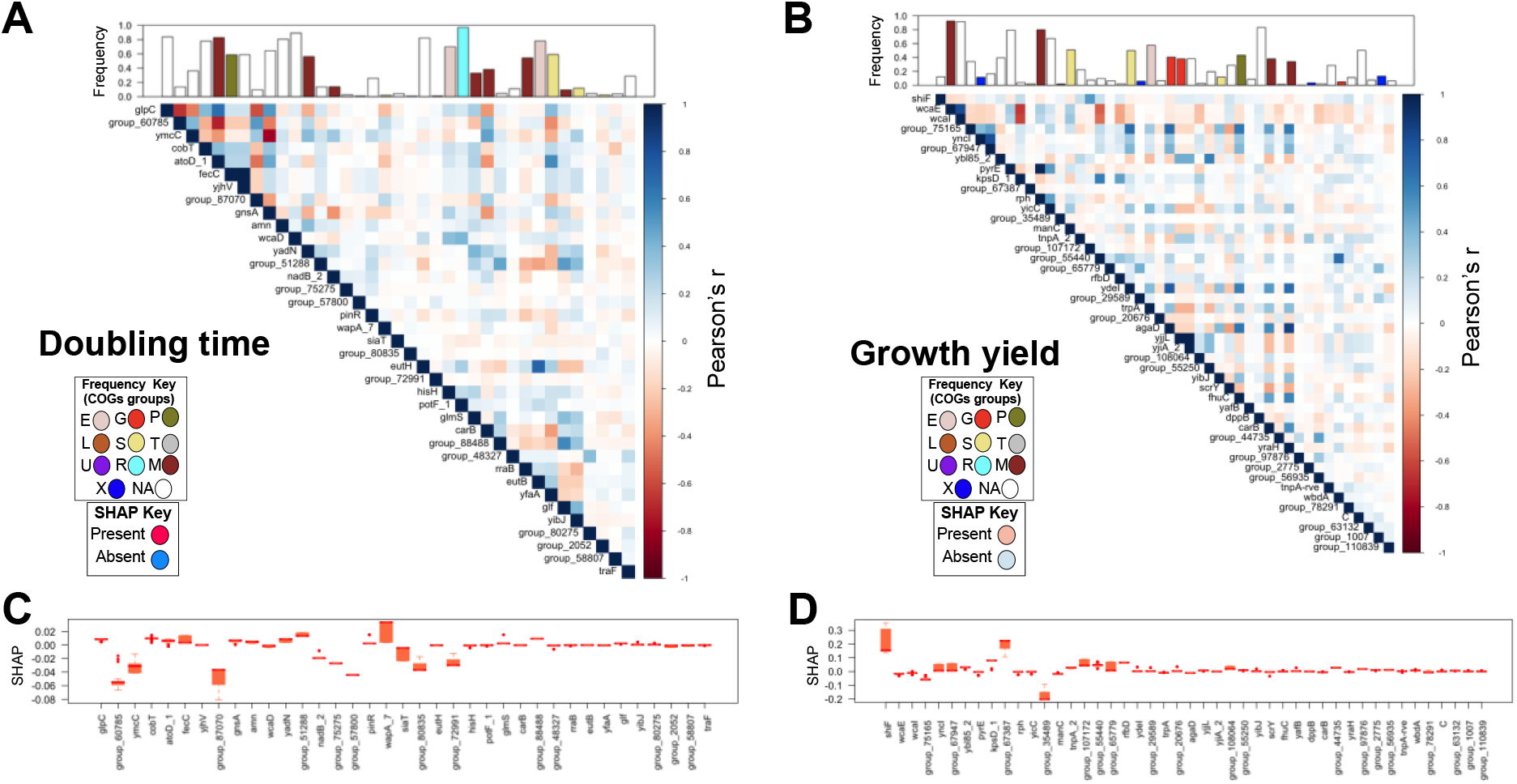
Feature importance analysis for the growth-related response features, i.e. A) growth yield and B) doubling time in the absence of antimicrobials. Features are sorted according to their average ranks across 20 prediction replicates. The bar-plots show the frequency of genes in the pan-genome. Colours represent the most prevalent COG functional groups (see Methods). The definitions of these groups are provided in http://www.ncbi.nlm.nih.gov/research/cog#. Genes *yjhv*-*fecC and kspD-rfbD* were found significantly linked with doubling time and growth yield after accounting for population structure, respectively (significance level p-value< 0.05). The matrix is pairwise association between presence of hits, where the colour density shows the strength of association and colors show the direction of the Pearson correlation. The box-plots show the SHAP value, i.e. the effect of the presence or absence of the genes on the response features, C) doubling time and D) growth yield.

We next extracted the gene presence and absence features that predicted growth at sub-inhibitory antimicrobial concentrations, and found between 23 and 250 accessory genes whose presence always were utilized by models to predict growth across 18 antimicrobial treatment conditions for growth yield and doubling time. The distribution of the majority of these genes were linked with the population structure, making the latter an unlikely cause for their association to antimicrobial resistances. Among the genes with phylogenetically reliable signals, we found many known AMR gene families (Table 1, Supplemental Table S2). The presence of *aacA-aphD* and *neo* (KAN) resistance, *cat* (CAM), *tet* (TET) and *dfrI* and *dfrV* (TRIM) consistently predicted a high cell yield in presence of the expected drug, whereas the presence of *aacA* and *aphD* (KAN) and *tet* (TET) also consistently predicted a short population doubling time in the corresponding condition (Table 1A and 1B). Our analysis also identified other gene families linked with resistance genes, including transposons linked with tetracycline resistance, *tnpA* (transposase) and tetracycline repressor gene *tetR*. Being carried on plasmids also containing antimicrobial resistance genes also explains why genes required for plasmid transmission and replication and phage or transposon factors predict growth in some antimicrobials, with e.g. the transposon *tnpA.* The *tnpA* is associated with Tn3 and has been reported in full or truncated forms downstream of diverse plasmid backgrounds containing *bla-CTX-M* and *bla-TEM* genes, respectively (43). A database search of AMR genes for similar genes identified ESBL, e.g. *bla-CTX-M* and *bla-TEM* genes, and resistance genes with overlapping regions with the inferred open reading frame for *tnpA* gene copies in the population.

**Table 1.**
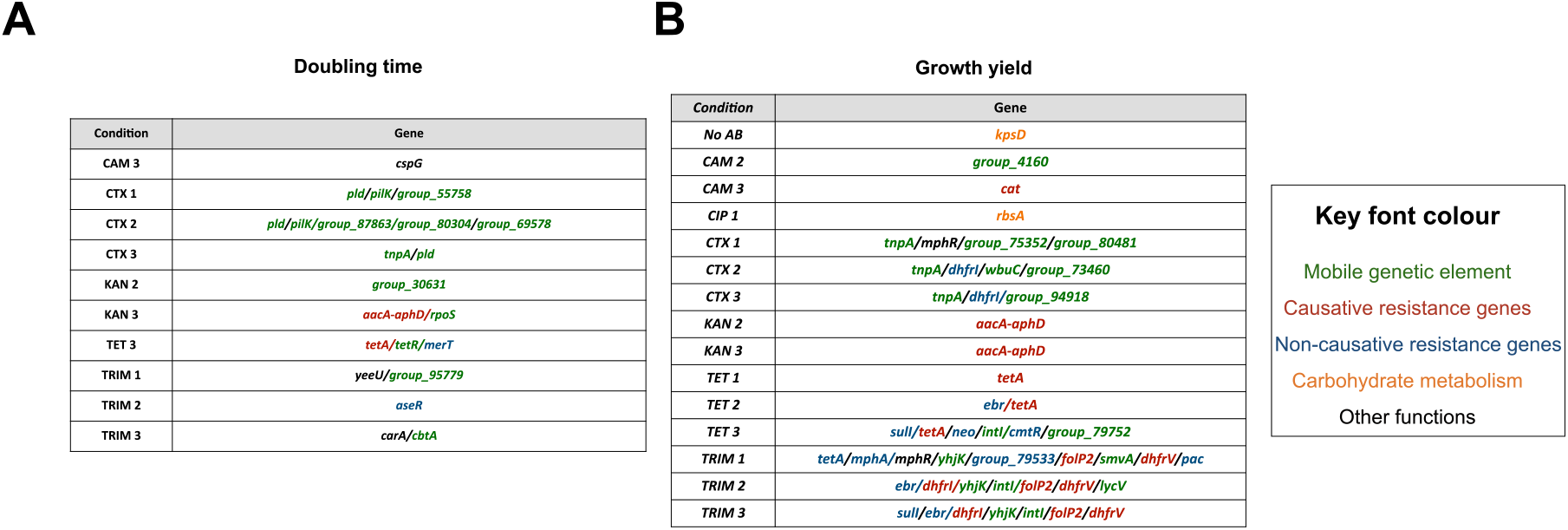
Predictive biomarkers for doubling time and growth yield, found significantly linked with the phenotype after accounting for population structure for different treatment conditions, i.e. drug type and concentrations (p-value cut-off <0.01). The full annotation of the genes is available in Supplemental Table S2.

The overlap between genomic features that were used to predict growth in different conditions was low (average Jaccard distance of 0.14 and 0.23 for growth yield and cell doubling time features, respectively), but among the few genes whose presence predicted tolerance to multiple antimicrobials, we found the well-known multidrug resistance genes, e.g. *neo* and *dfrI,* reflecting the extensive cross resistance and cooccurrence of multiple resistance genes (Table 1). We used the SHAP values to quantify the impact of predictors on the growth of each strain and first estimated the contributions of known resistance genes, first individually (Figure 4). The presence of the of the tetracycline efflux pumps gene family *tet* improved the growth yield with on average 15% of the total population range and reduced the cell doubling time with on average 11% of the total population range (Figure 5). The impact of the presence of these genes is already noticeable at low tetracycline concentrations.

**Figure 4.**
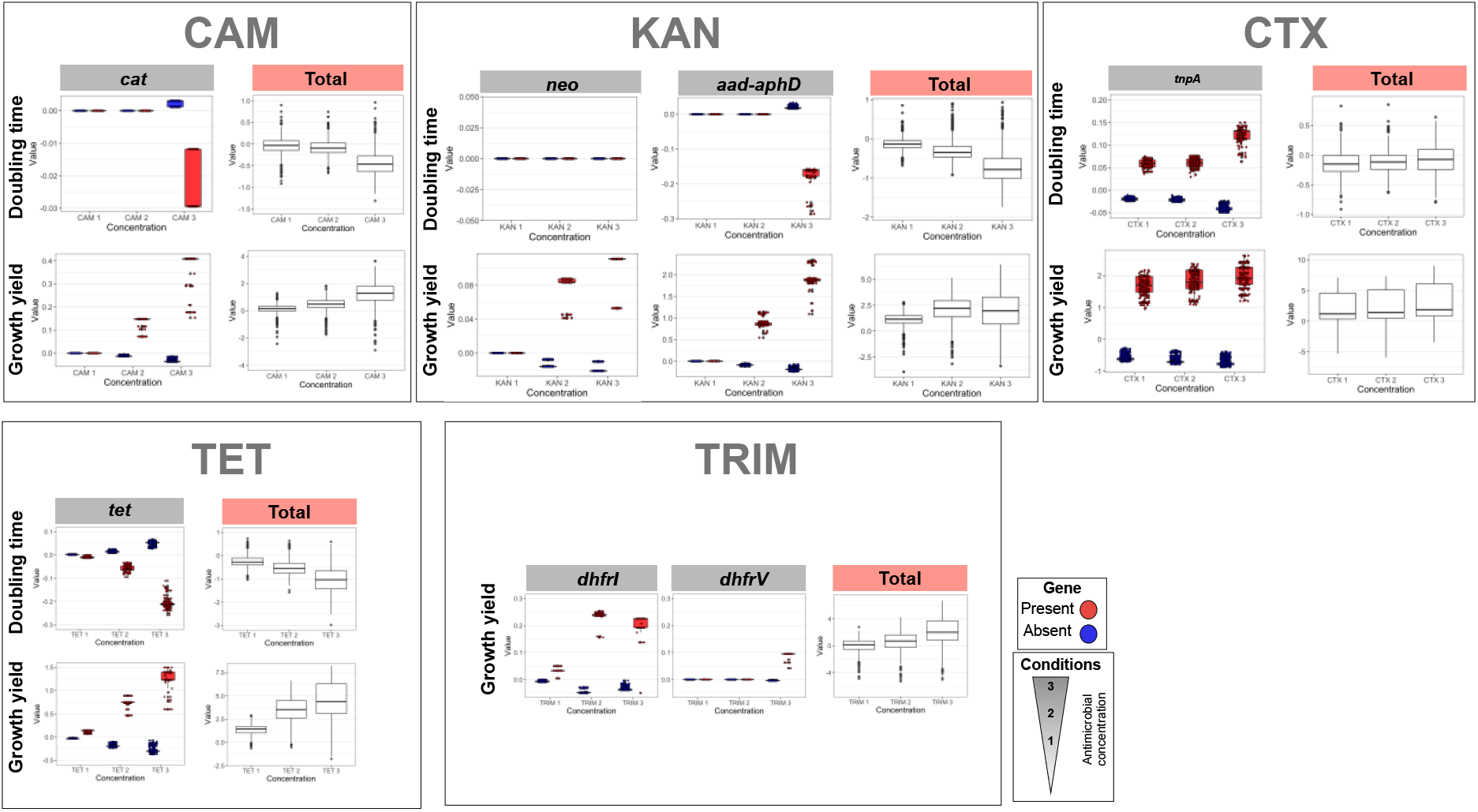
The contribution of the presence of known resistance genes and *tnpA* gene to the prediction of growth-related features, i.e. growth yield and doubling time, as measured by SHAP values. The *tet* gene family includes variants *tetA, tetD* and *tetC* of the genes in the CARD database. Numbers above each plot shows the frequency of the gene. The *tnpA* gene was found linked with ESBL genes, i.e. *bla-CTX* and *bla-TEM*. Boxplots in red and blue correspond to distribution of the effect of the presence and absence of features on growth-related features in each sample, respectively. Boxplots in white show the population distribution of growth-related features for the purpose of comparing the effect of the genes.

The two dihydrofolate reductase genes, encoded by *dfrI* and *dfrV*, both improved the growth yield under high trimethoprim exposure, with individual contributions of 3% and 1% of the total population range, respectively, but had no effect on cell doubling time. We examined fitness effects (growth yield increase) of different combination of *dfrI* and *dfrV* at high concentrations. This corresponded to a single-peaked fitness landscape, with *dfrI* contributing much more to resistance than *dfrV* (Figure S5). Thus, lineages only carrying *dfrI* achieved 99% of the cell yield of those containing both *dfrI* and *dfrV*, as compared to 3% for lineages containing *dfrV* alone (Figure S5). This reflects a slight negative epistasis, consistent with the diminishing return of beneficial genes and variants reported in *E. coli* lab strains (44). The presence of chloramphenicol acetyltransferase gene, *cat*, improved the cell yield by 0, 5% an 12% of the total population range in low, moderate and high concentrations of chloramphenicol but only marginally reduced the cell doubling time (0.02% in the highest concentrations) (Figure 4). Similarly, the KAN resistance aminoglycoside phosphotransferase genes family *aacA-aphD* individually improved the cell yield with 0, 12% and 31% at low, moderate and high kanamycin concentrations, but reduced the cell doubling time only slightly and only at the highest concentration (Figure 5). Similar to *aacA-aphD,* the presence of amino 3′-glycosyl phosphotransferase (*neo*) gene, increased the growth yield at intermediate and high concentration concentrations but at only at 5-10% of the effect of *aacA-aphD* (Figure 4). Furthermore, the gene does not seem to affect doubling time. The presence of the transposon *tnpA*, reflecting the impact of the linked *bla-CTX-M* and *bla-TEM* genes, was associated with an increased growth yield with 10-13% of the total population range across antimicrobial concentrations (Figure 4), but had a consistently detrimental impact on the cell doubling time. Overall, our predictions captured much of the AMR biology and made both qualitative and quantitative sense, with known AMR resistance genes contributing mostly to growth yield rather than doubling times.

While the contribution of gene presence-absence variation to AMR is relatively well understood, the contribution of SNPs is less well explored. We therefore next considered the effect of 39 synonymous, 31 non-synonymous and 46 intergenic significant variants (see Methods) on the predicted cell doubling time and cell yield (Figure S6A, Supplemental Table S3). The *gnd* gene, which is part of the pentose-phosphate carbohydrate catabolism pathway, harboured the highest number of non-synonymous variants consistently used by predictions (Figure S6B). The gene is highly polymorphic within *E. coli*, owing to inter-strain transfer and recombination. Interestingly, the gene is situated close to the O antigen determining *rfbD* region in the *rfb* locus, reported above, suggesting the implication of the region in bacterial grwoth (45). Other genes with more than one variant used by models included genes encoding autotransporters and promoting biofilm formation (*yeeJ*) (46), cell division inhibition (*dicB*) (47), flagellar components (*filF*) and *kdpE* gene, encoding an activator of genes involved in a high-affinity potassium uptake system (48) (Figure S6B). We found two A84P and D87G variants in the Quinolone resistance-determining region of *gyrA* (49, 50), predicted to confer resistance to CIP concentrations to contribute to the cell yield during subMIC concentrations of CTX, TRIM and KAN exposure. Given that our collection includes multidrug resistant clinical strains, this observation indicates how extensive cross-resistance between strains cause resistance variants for one antimicrobial to be predictive for other antimicrobials. We also explored the genes containing non-synonymous sites that were caused by more than one type of base alteration in the population (Methods), and found three sites in the *gnd* gene and two in the O-antigen capsule biosynthesis *wcaM* gene (Supplemental Table S3). The presence of the membrane, capsule production and regulatory genes in the SNP and gene-based analysis suggest a key role for these genes in growth or stress response to antimicrobial treatment, although the causative link needs to be demonstrated in forward genetic experiments.

Pan-genomes show a wide range of sizes across species, with accessory genome comprising 16% to 97% of total for well-sampled genomes across bacterial species (51). We therefore examined whether variation in pan-genome sizes would affect prediction, which informs on whether our models would be predictive in other bacterial species. We therefore repeated the prediction of cell yields and cell doubling time at subMIC antimicrobial concentrations for simulated pan-genomes with increasing accessory genome size and for increasing population size. As anticipated, the prediction error and the extent of overfitting, i.e. the difference between the error rate for training and test dataset, monotonically decreased as population size increased (Figure S7A), although a level of overfitting existed at all population sizes tested. The monotonic effect of decreasing pan-genome size, and potentially the background noise from uncorrelated genes, on improving prediction could also be seen. When we decreased the pan-genome size by 100 times, the true detection rate of the causative genes remained in the same range (49%,46% and 48% for gene acquisition values of 10^−7^, 10^−8^ and 10^−9^, corresponding to accessory genome size of 77%, 23% and 1% of pangenome, respectively), however an increase in sample size improved the detection rate (48%, 59% and 61% for sample size of 200, 400 and 600, respectively) (Figure S7B). Moreover, the correlation between predicted and actual values on the test dataset was positive for all simulated datasets for selective advantage as low as 0.5 for the causative gene in all scenarios for population and pan-genome sizes. This value is greater than the estimated selective advantage for the resistance gene under 15 out of 19 treatment conditions for growth yields in Figure S4A (see Methods). These results show that the predictive model remains applicable to a broad range of scenarios for pan-genome evolution.

## Discussion

In this study, we adopted a reverse-genetic approach and applied three different machine learning models to predict bacterial growth and doubling time from genomic data under a range of growth conditions in natural *E. coli* strains. We focused on interpreting features of machine learning models to advance a mechanistic understanding of the genetic repertoire for growth under antimicrobial treatment.

We concluded that tree-based models were superior to standard fully connected deep neural network models for AMR prediction from genomic variants, confirming previous successful applications of ensemble-boosting classifier and regressor methods for inferring AMR phenotype and pathogenicity from genomic data in Gram-negative strains (10, 12). This recurring pattern may be due to the nature of these datasets, in which features are individually meaningful and a strong multiscale structure is absent. For the purposes of routine and effective clinical use, these models still require interpretability, as the mechanisms and patterns that the model uncovers are important for practical applications. Our findings revealed how the interpretability of features could provide an understanding of the model when local explanations of each prediction were combined. Using this approach, we measured the marginal effect of biomarkers on prediction across different combinations of allelic status. Due to our modest sample size, we limited our analysis to known resistance determinants. However, the rapid increase in genomic data will allow future studies to extend the approach to infer fitness effects for a large number of genomic variants with moderate-to-weak effects. Such analysis not only allows comparing the relative importance of resistance genes across conditions but assessing the average contribution of a gene across many genomic backgrounds, which is infeasible and laborious in the lab. The data-driven approach is also superior to experimentally assessing fitness effects of genes in individual lab strains over the course of the evolution, since this approach accounts for various genetic backgrounds, through which the gene is passed. This is attained by computing the average fitness effect and the extent of variation around that average, which becomes doubly valuable when considering the average effect in presence/ absence of other genes with an effect.

Antimicrobial resistance research has been largely focused on clinical strains and therapeutic antimicrobial concentrations. Therefore, the contribution of sub-inhibitory antimicrobial traces, released as a result of anthropogenic interventions in the environment, to the rise of resistance has not been sufficiently studied (52, 53). Prior evidence suggests that many types of plasmid resistance do not emerge *de novo* during treatment (54). Thus, a patient or animal is either infected with the susceptible bacteria, without any resistance developing during treatment, or they are infected with the resistant strain and the antimicrobial treatment mainly causes an enrichment of a pre-existing resistance gene or mutation. Our results demonstrated that the impact of resistance genes is detected at sub-inhibitory concentrations. Therefore, the fitness effects appear to outweigh the fitness cost at low antimicrobial concentrations. This further supports the idea that the ability of most resistance genes to confer high-level resistance at a low fitness cost shields the selective dynamics of mutants at low drug concentrations, which leads to the selection and fixation of resistance variants (16). This finding calls for the extension of the selection window in resistance stewardship programs to include sub-inhibitory antimicrobial concentrations, specifically the minimum selective concentration (MSC) (53, 55). This applies particularly to environmental sites, where the selection for antimicrobial levels might be ongoing.

Despite attaining a strong (56) average correlation of 0.63 and 0.47 between predictions for growth yield and population doubling time, respectively, there remains a large gap to perfect prediction. We do not attribute the low heritability to gene expression variation. The reason is that the variation, irrespective of the nature of the variation, i.e. environmental or stochastic, is expected to be captured in measurement errors, which we have accounted for in our models. A number of other factors may explain the low heritability. Some studies suggested a possible role for sequence independent determinants of resistance, e.g. epigenetics, in the development of resistance to the antimicrobial, in particular at sub-inhibitory concentrations of antimicrobials (57–59). Another possible explanation is the role of rare variants or variants with weak fitness effect remaining to be discovered (60). These could be either common alleles with moderate effects or rare alleles with large effects. The latter has been reported for lab-colony pools, where around 10% of resistance to rifampicin was caused by numerous rare mutations (61). Furthermore, low-effect mutations different from classic high-effect drug resistance have been identified to drive streptomycin resistance at sub-MIC levels in *Salmonella enterica* (62). Hence, capturing the information of rare genetic variants, including variants types caused by genomic rearrangements and insertion sequence movements [e.g. (63)], for the genome-based prediction of resistance remains a challenge (64). The total heritability may also be increased by genetic interactions (60), at inter- or intra-gene levels (65), which limits the genome-based AMR prediction accuracy in small-to-moderate size genomic datasets (64). Besides limitations in prediction, our results also showed the extent to which cross-resistance, lineage and linkage may constrain the application of ensemble models, especially when the aim is to identify causative biomarkers. The lineage association for predictive features makes the robust identification of a universal biomarker gene complicated, as the performance of the trained machine-learning model will be sensitive to the lineages included in the training dataset. Therefore, a comprehensive and large-scale genomic sample is required to mitigate the above effects.

Deriving a genotype-phenotype map for *E. coli* strains from diverse sources has important implications for understanding and predicting the dynamics of the population on epidemiological timescales and across environmental and clinical sites. Moreover, such genotype-phenotype map may directly inform on potential novel targets to prevent the spread of clinically relevant strains in natural populations. The successful application of machine-learning methods provides the motivation for these methods to be employed in future studies to predict other clinically relevant traits, such as transmissibility, host-preference and horizontal gene transfer rate. These endeavours would meaningfully improve infectious disease diagnostics.

---

**Supplemental Figure S1.**
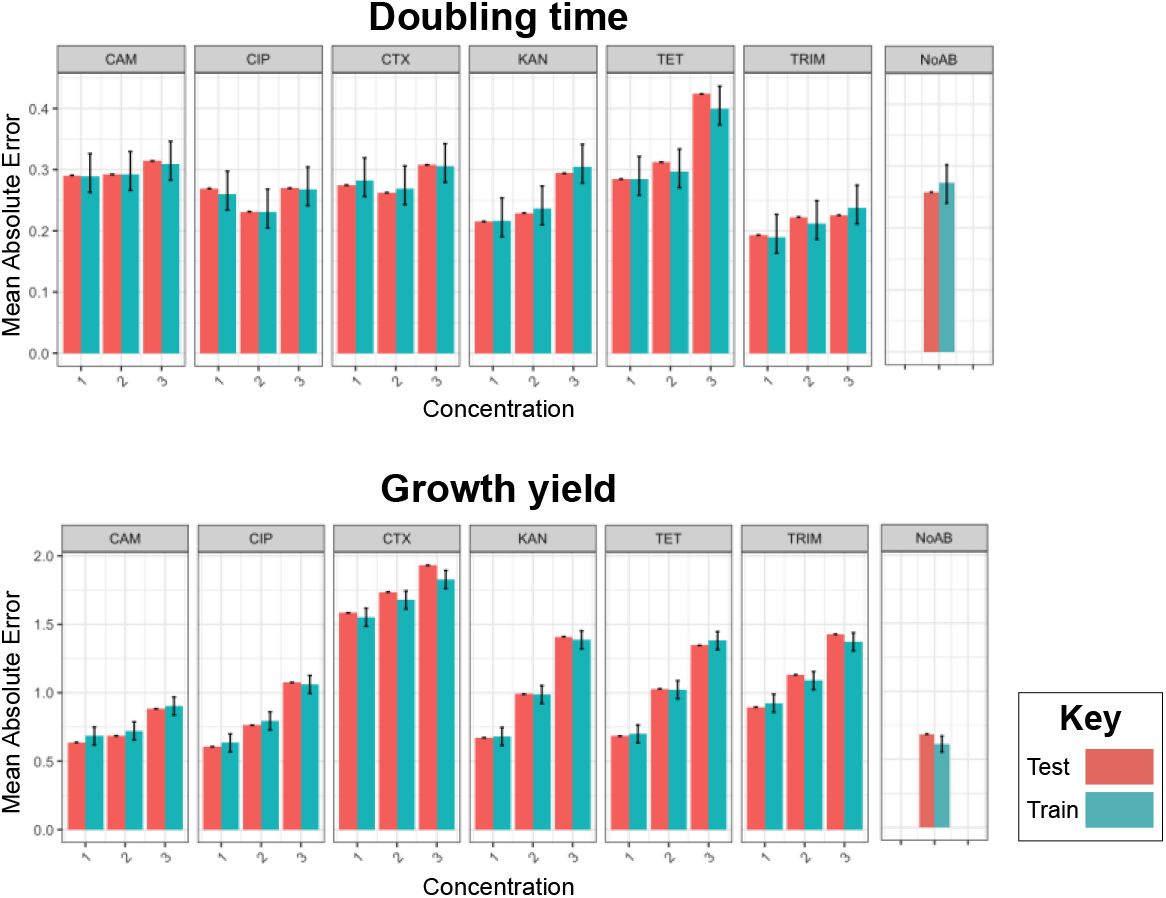
The Mean Absolute Error (MAE) values for training and test data for the tuned gradient regressor model. Error bars shows 95% confidence interval.

**Supplemental Figure S2.**
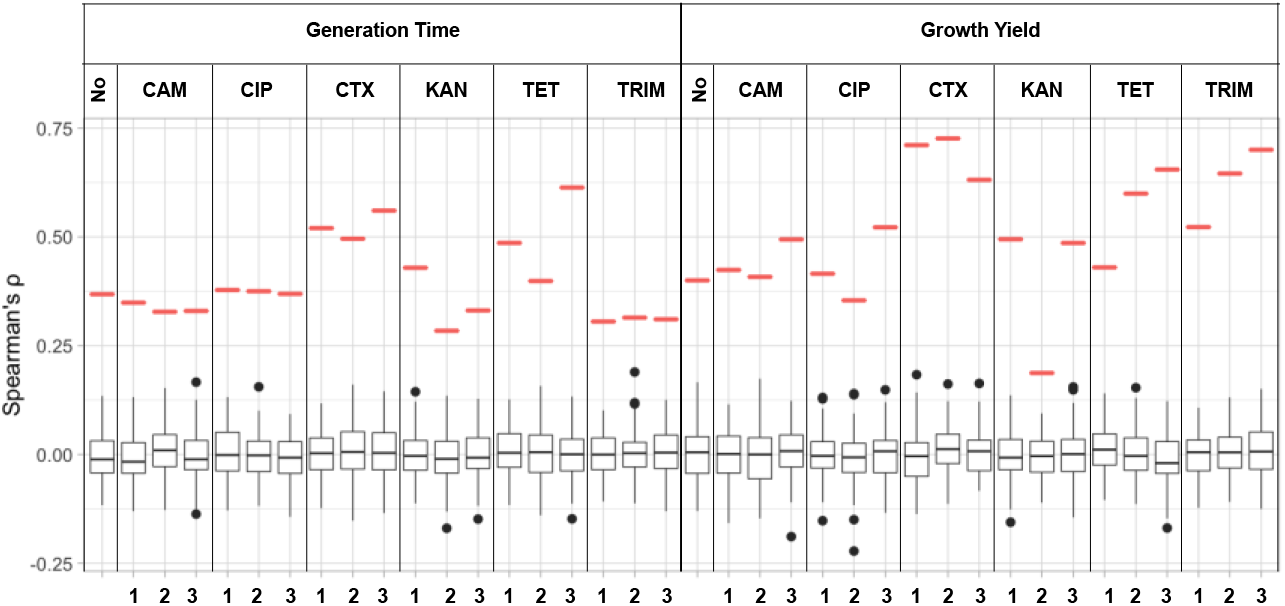
The significance of results for the best performing tuned gradient regressor model in Figure 1. The boxplots show the distribution of Spearman correlation coefficients for 100 bootstrapped sample sets from the original data.

**Supplemental Figure S3.**
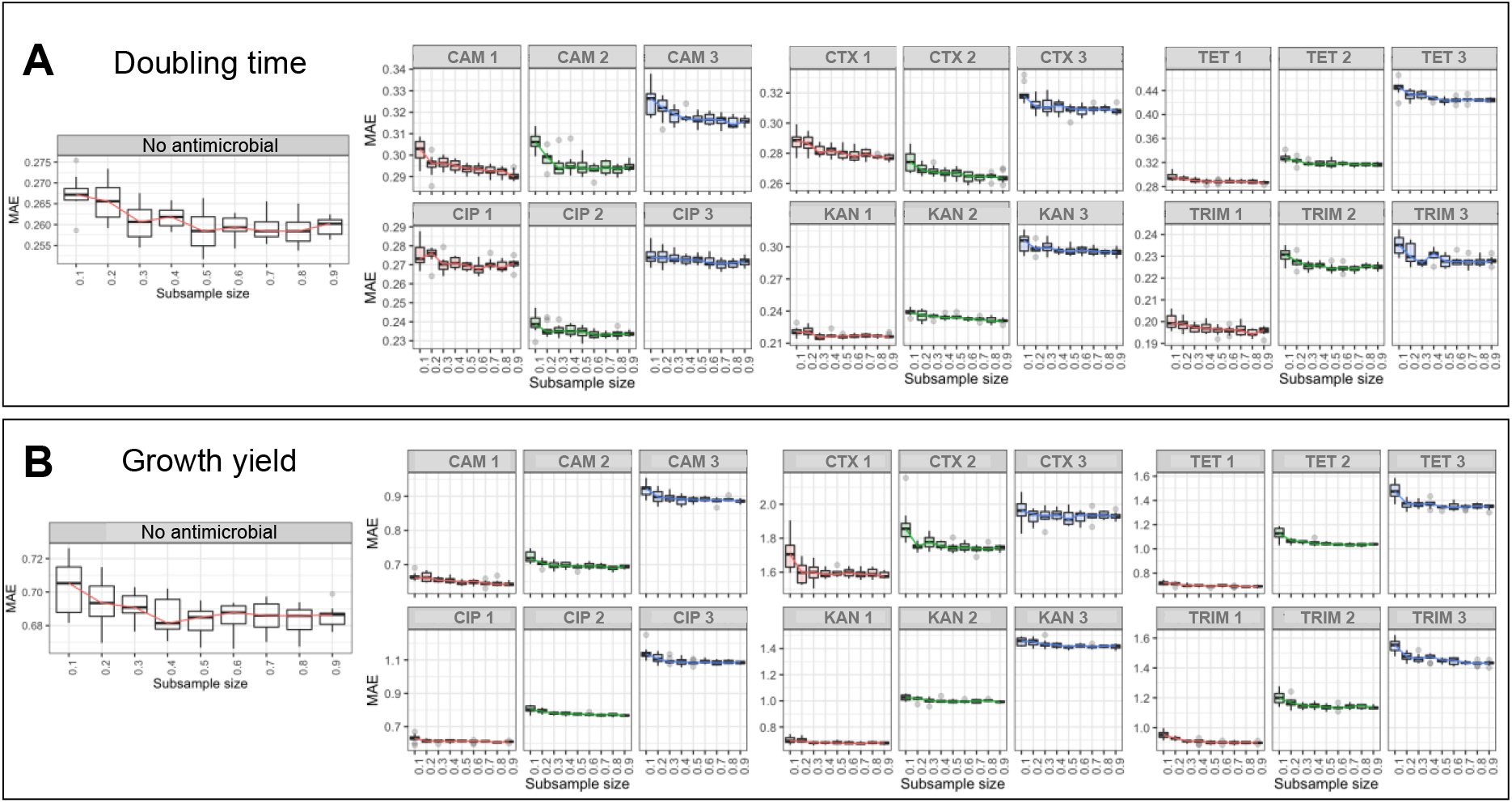
The Mean Absolute Error (MAE) values for training and test data for the tuned gradient regressor model. Each box shows the results for 100 randomly generated subsamples.

**Supplemental Figure S4.**
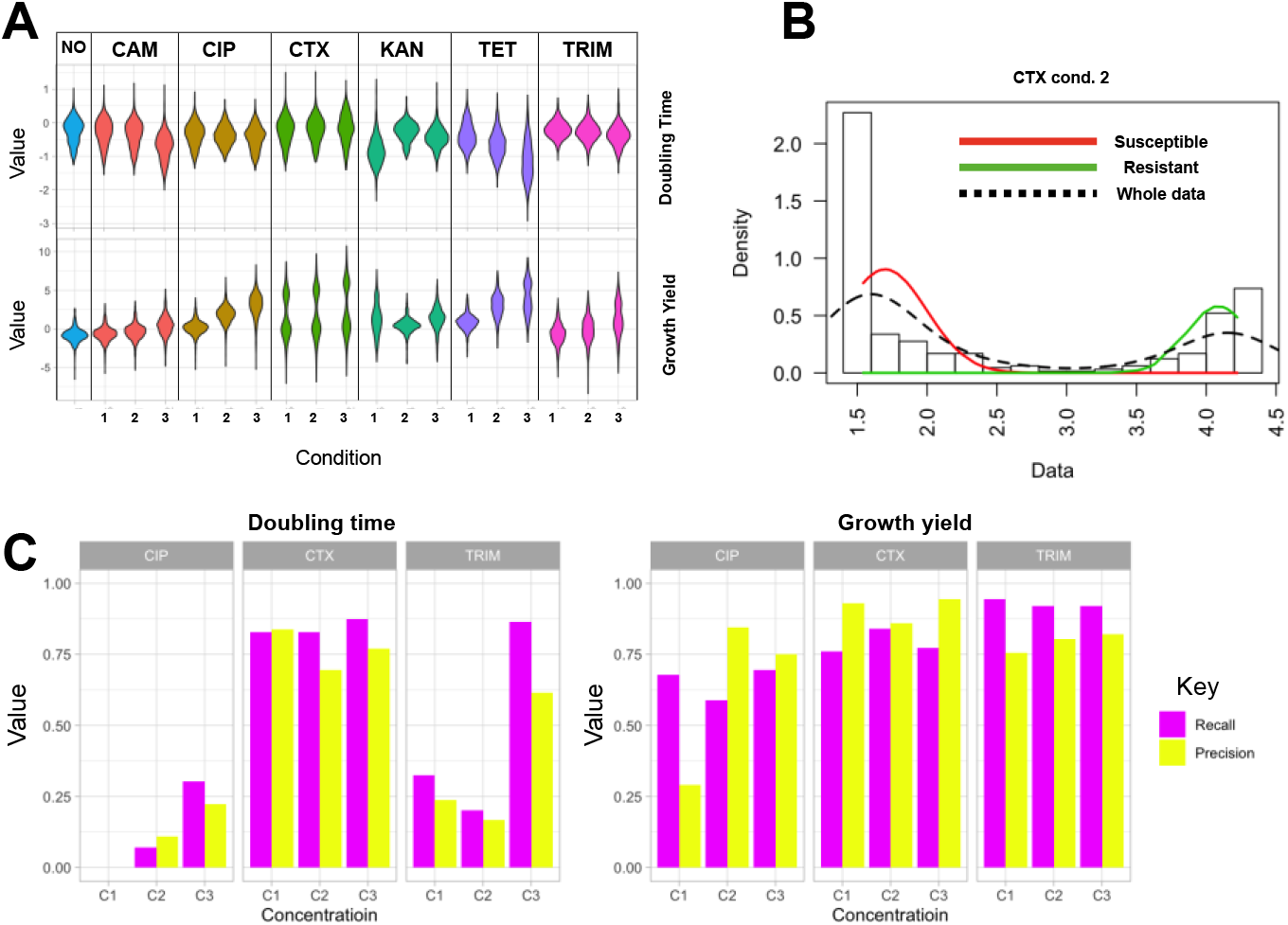
A) distribution of fitness assay data for doubling time and growth yield. B) An example of the predicted values for the growth yield for cefotaxime under the third condition. The dotted line shows the fitted curve for a bimodal distribution. The red and green lines show the inferred underlying normal distributions. C) Prediction of the resistant phenotypes using labels inferred from the predicted bimodal distributions for three antimicrobials. The purple and yellow bars show recall and precision for the resistant phenotype, respectively.

**Supplemental Figure S5.**
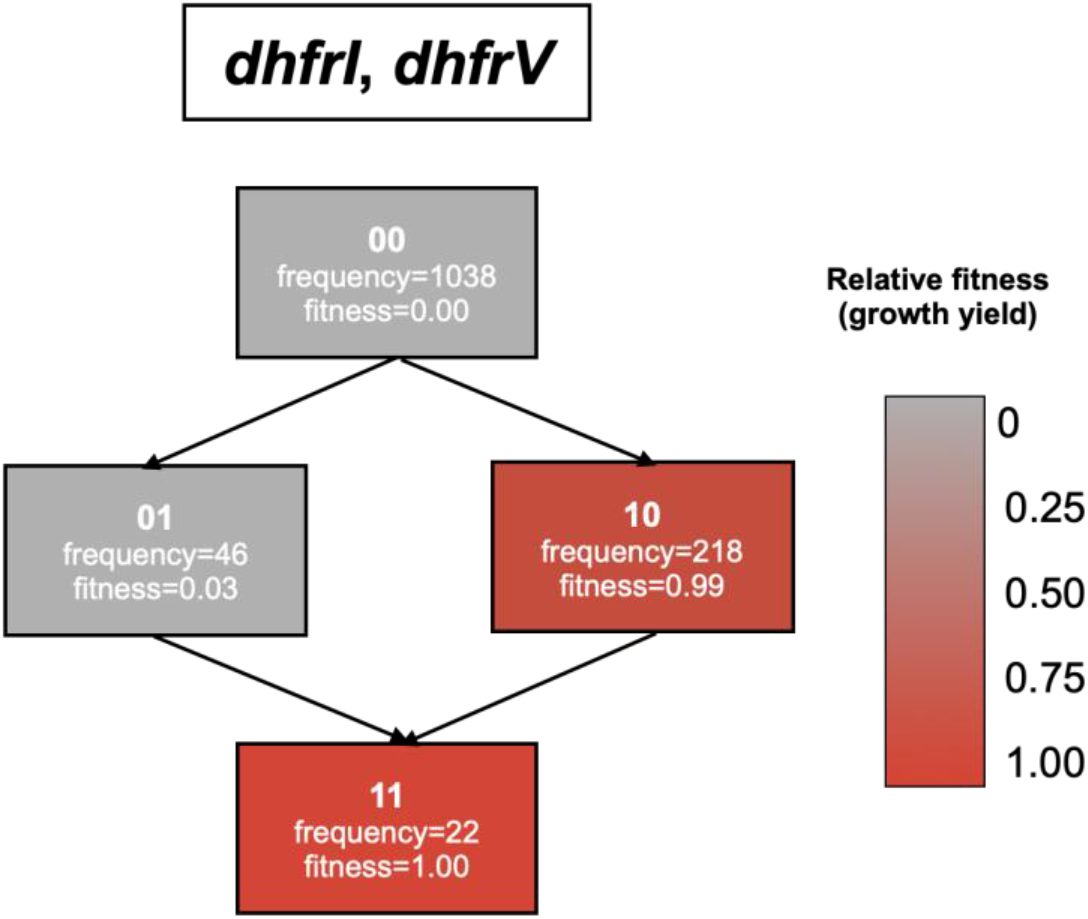
Fitness landscape of growth yields reconstructed from SHAP values for resistance genes against trimethoprim at the highest concentrations. The 1 and 0 signs represent the presence and absence of the genes in the order shown above the figures, respectively. Colours correspond to the mean SHAP values the specified combinations of resistance genes across all strains.

**Supplemental Figure S6.**
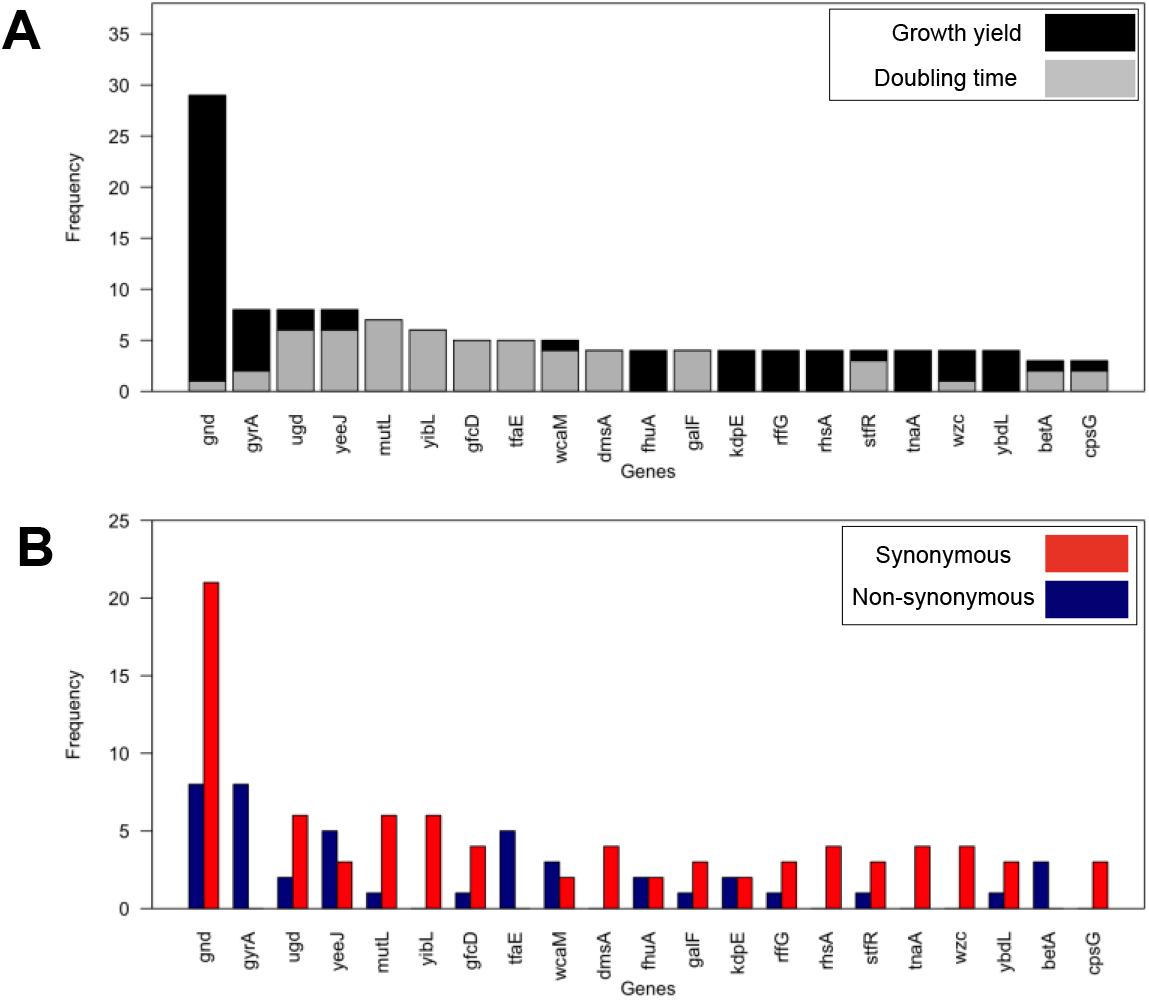
List of genes with predictive SNP biomarkers that were significantly linked (p-value from ANOVA test< 0.05) growth conditions measures in 38 conditions for doubling time and growth yield in the absence and presence and 3 conditions for six antimicrobials. The complete list of mutations is provided in Supplemental Table S3.

**Supplemental Figure S7.**
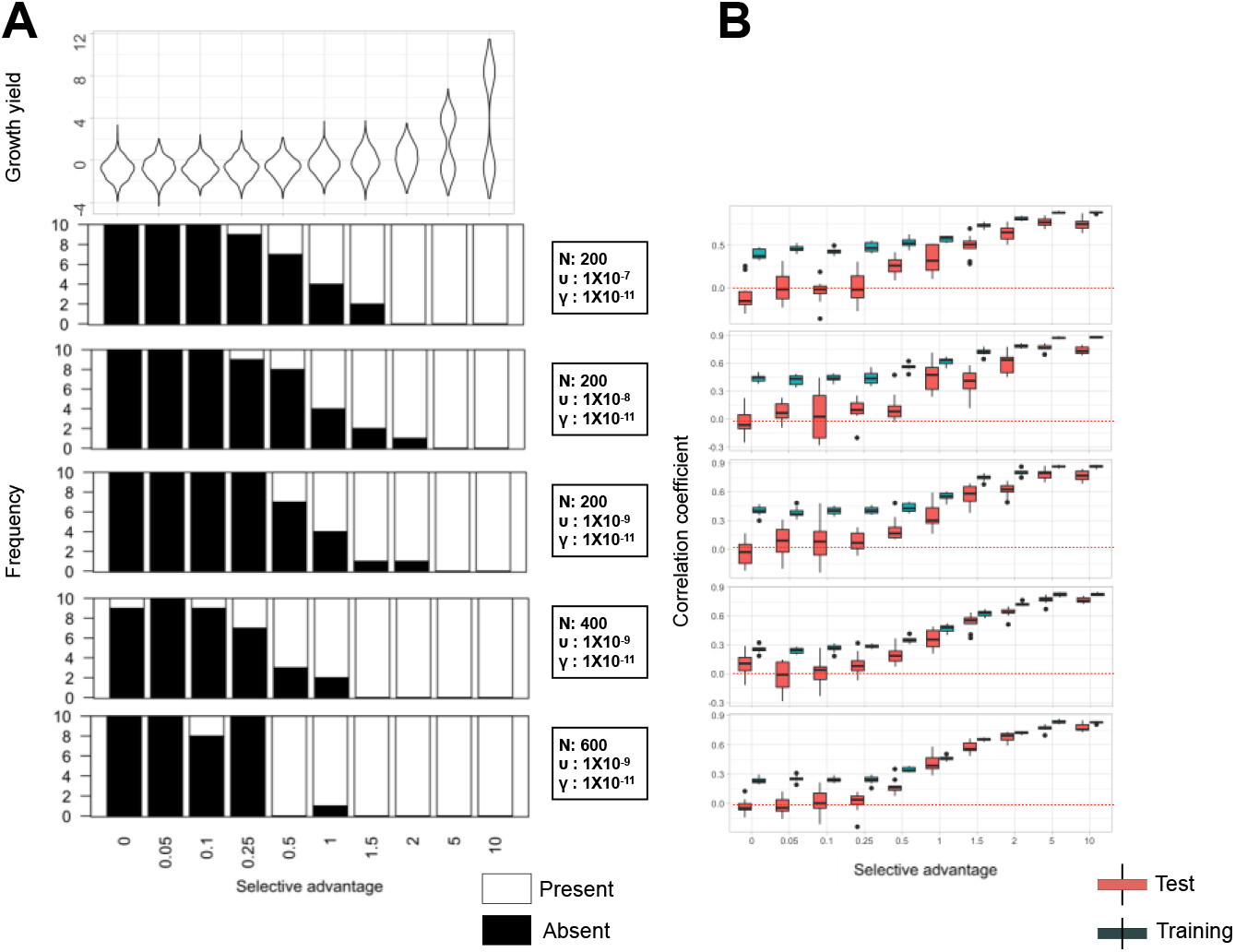
The frequency of correct prediction of the causative gene for simulated pan-genomes and increasing value for the selective advantage of 10 causative genes. White bars correspond to cases where the causative gene is correctly identified by the tuned gradient boosted regressor model in a test data set. Further details are provided in Methods. Signs N, υ and γ signs show population size, rate of gene acquisition and rate of gene loss for reconstructed pan-genomes. We used gradient boosted regressor model with 100 iterations and default parameters.

## Acknowledgment

This work was partially funded by a grant from the Centre for Antibiotic Resistance Research (CARe) at the University of Gothenburg to AF and grant number 2016-06503 from the Joint Programming Initiative on Antimicrobial Resistance (JPIAMR) to JW and AF. This work was in part supported by the Academy of Finland (grant 313270 to VM). LP was supported by Wellcome, Estonian Research Council (IUT34-4), Estonian Centre of Excellence in IT (EXCITE) (TK148). DM was supported by the Joint Programming Initiative on Antimicrobial Resistance (JPIAMR) via MRC grant MR/R004501/1. The funders had no role in study design, data collection and analysis, decision to publish, or preparation of the manuscript. TYD is participated in the Erasmus and Mobility for higher education project, sponsored by the British Council (2018-1-TR01-KA103-049917). The Culture collection University of Gothenburg (CCUG), Daniel Jaén-Luchoro, Christina Åhern, Nahid Karami, National collection of type cultures (NCTC), Carl-Fredrik Flach, Jan Michiels and Marco Galardini are gratefully acknowledged for strains.

## Supplemental Tables

Supplemental Table S1: Samples specification and accession Supplemental

Table S2: List of significant accessory genes Supplemental

Table S3: Definition of significant SNPs

